# Temporal regulation of nicotinic acetylcholine receptor subunits supports central cholinergic synapse development

**DOI:** 10.1101/790659

**Authors:** Justin S. Rosenthal, Jun Yin, Caixia Long, Emma Spillman, Chengyu Sheng, Quan Yuan

## Abstract

Construction and maturation of the postsynaptic apparatus are crucial for synapse and dendrite development. The fundamental mechanisms underlying these processes are most often studied in glutamatergic central synapses in vertebrates. Whether the same principles apply to excitatory cholinergic synapses in the insect central nervous system (CNS) is not known. To address this question, we investigated *Drosophila* ventral lateral neurons (LNvs) and identified nAchRα1 (Dα1) and nAchRα6 (Dα6) as the main functional nicotinic acetylcholine receptor (nAchR) subunits in these cells. With morphological and calcium imaging studies, we demonstrated their distinct roles in supporting dendrite morphogenesis and synaptic transmission. Furthermore, our analyses revealed a transcriptional upregulation of Dα1 and downregulation of Dα6 during larval development, indicating a close association between the temporal regulation of nAchR subunits and synapse maturation. Together, our findings show transcriptional regulation of nAchR composition is a core element of developmental and activity-dependent regulation of central cholinergic synapses.

## Introduction

The postsynaptic compartment of a chemical synapse is specialized to receive neurotransmitter signals and translate them into electrical and chemical changes in the postsynaptic cell. Its establishment and modification are critical for the development and plasticity of synapses and dendritic arbors and are regulated by the coordinated efforts of genetic programs and external influences. The structural and functional organization of postsynaptic specification has been investigated extensively in the excitatory glutamatergic synapse of the vertebrate central nervous system (CNS), where the NMDA and AMPA-type of ionotropic glutamate receptors (iGluRs) are concentrated at the postsynaptic densities (PSD), a network composed of scaffolding proteins, cell adhesion molecules and signaling proteins (Frank and Grant, 2017; Sheng and Kim, 2011). During development, NMDA receptor-mediated calcium signaling induces dendrite growth and synapse partner selection (Lohmann and Bonhoeffer, 2008; Sin et al., 2002), while the changing AMPA/NMDA ratio regulates synapse strength and contributes to the physiological maturation of the glutamatergic synapse (Ben-Ari et al., 1997; Pratt et al., 2016; Wu et al., 1996).

In contrast to the well-established role of glutamate receptor signaling in sculpting the developing vertebrate brain, it is unclear whether neurotransmission mediated by acetylcholine (Ach) is also a major effector of CNS development and plasticity. Although both ionotropic nicotinic (nAchR) and metabotropic acetylcholine receptor (mAchR) signaling contribute immensely to animal behavior and are identified as the neural substrates for a multitude of neuropsychiatric disorders in humans, our scope of understanding their neuronal function is not yet comparable to that of the central glutamatergic synapses (Ballinger et al., 2016; Dineley et al., 2015; Rosenberg et al., 2002). In the mammalian brain, Ach is studied primarily as a neuromodulator; extra-synaptic nAchRs modify axonal release and/or receptor sensitivity to glutamate, GABA and other neurotransmitters, and are expressed in distinct regions of the mammalian brain such as the ventral tegmental area and nucleus accumbens (Ballinger et al., 2016; Picciotto et al., 2012). On the other hand, in the vertebrate neuromuscular junction (NMJ) and insect CNS, Ach is known for mediating fast, ionotropic synaptic transmission (Gu and O’Dowd, 2006; Gundelfinger, 1992; Picciotto et al., 2012; Wu et al., 2010).

The *Drosophila* model provides an excellent system to expand this field of study, not only because many central synapses in insects use Ach as the main excitatory neurotransmitter, but also because of the diversity of nAchR subunits in the *Drosophila* genome. However, the molecular composition and regulatory mechanisms of the *Drosophila* postsynaptic nAchR complex remain largely unknown (Campusano et al., 2007; Dupuis et al., 2012; Gu and O’Dowd, 2006; Jones and Sattelle, 2010). A few nAchR genes have been ascribed behavioral functions, including the *Drosophila* nicotinic receptor 7 (Dα7), which is expressed in giant fibers and mediates the escape reflex, and Dα1 and Dα3, which help regulate complex behaviors such as courtship and sleep (Fayyazuddin et al., 2006; Shi et al., 2014; Somers et al., 2018; Wu et al., 2014), but information about the cellular and physiological functions of nAchRs remains limited. Nonetheless, the complex yet critical roles of nAchRs in the development and organization of cholinergic synapses have been implicated. Previous studies demonstrated the synaptic distribution of Dα6 and Dα7 subunits and the synaptic transmission mediated by nAchRs in a memory-relevant mushroom body output synapse (Barnstedt et al., 2016; Kremer et al., 2010; Nakayama et al., 2016).

To understand whether and how nAchR signaling contributes to the development and plasticity of the central cholinergic synapse, we performed genetic studies in *Drosophila* larval ventral lateral neurons (LNvs), a group of visual projection neurons receiving nAchR-mediated excitatory cholinergic transmission from larval photoreceptors (Wegener et al., 2004; Yuan et al., 2011). Notably, synapse formation and dendrite development in LNvs are both strongly influenced by changes in visual input, and thus provide a unique system for genetic dissections of activity-dependent regulation of synapse and dendrite development (Sheng et al., 2018; Yuan et al., 2011).

While the native compositions of pentameric nAchR ion channels in *Drosophila* central neurons are generally not known, through cell-specific RNA-seq analyses, morphological screens and calcium imaging studies, we identify Dα1 and Dα6 as the main functional nAchR subunits in LNvs and demonstrate their distinct roles in supporting synaptic transmission and dendrite morphogenesis. Furthermore, our analyses reveal a transcriptional upregulation of Dα1 and downregulation of Dα6 during larval development, indicating an association between the developmental switching of nAchR subunits and the maturation of central cholinergic synapses. Together, our studies of phenotype and temporal regulation show the two nAchR subunits Dα1 and Dα6 playing distinct functional roles during LNv development: *Dα6* is required in the early synaptogenesis period, whereas *Dα1* expression is potentiated at a later stage to fine-tune the physiological properties of LNvs and support synaptic transmission in a mature cholinergic synapse.

## Results

### Sequence alignment reveals distinct subgroups among the *Drosophila* nAchR subunits

The genome of *Drosophila melanogaster* contains 10 nAchR subunit-encoding genes, which, like those of other insect nAchR families, are fewer in number than the 16 found in humans and 27 in *C. elegans* (Jones and Sattelle, 2010; Schafer, 2002). Seven genes encode the alpha subunits (Dα1-7) and three encode the beta subunits (Dβ1-3) of nAchRs. Although all subunits are characterized by similar sequence motifs, such as those encoding the 15-amino acid Cysteine-loop (Cys-loop) and four transmembrane domains, the homology alignments incorporating human nAchR subunits uncovered phylogenetic relationships within the family (Supplementary Figure 1). We found that subunits Dα5-Dα7 are clustered with the human nAchRα7 (*CHRNA7)*, which is known for its ability to form homomeric pentamers and its high conductance of Ca^2+^ ions (Albuquerque et al., 2009; Dani, 2015). The other *Drosophila* subunits, *Dα1-Dα4, Dβ1* and *Dβ2*, are dispersed among the remaining human nAchRα (*CHRNA)* and nAchRβ (*CHRNB)* genes, as well as the non-neuronal γ, δ and ε subunits. The *Dβ3* subunit appears to be the outgroup in this analysis, consistent with the identification of species-specific nAchR genes in other insects which are highly divergent from those of their own genome (Jones et al., 2007).

### *Dα1* and *Dα6* are identified as candidate genes regulating synapse and dendrite development in larval LNvs

LNv dendrites receive cholinergic inputs from larval photoreceptors through the nAchRs, suggesting cholinergic receptor clusters serve as key components of LNvs’ postsynaptic specializations (Wegener et al., 2004; Yuan et al., 2011). To identify the specific nAchR subunits expressed in larval LNvs, we first analyzed the relative abundance of transcripts for all AchR subunits using cell-specific RNA-seq analyses (Yin et al., 2018) (Figure 1A). At the late 3^rd^ instar stage, most of the AchR subunits were detected in the LNv transcriptome, except for Dβ3 (Figure 1B). Notably, LNvs expressed high levels of Dα1 with minimal Dα7 expression, a pattern different from that found in cholinergic synapses of the adult olfactory circuit, visual system and giant fibers where the Dα7 subunit is enriched (Fayyazuddin et al., 2006; Kremer et al., 2010; Leiss et al., 2009).

**Figure 1.**
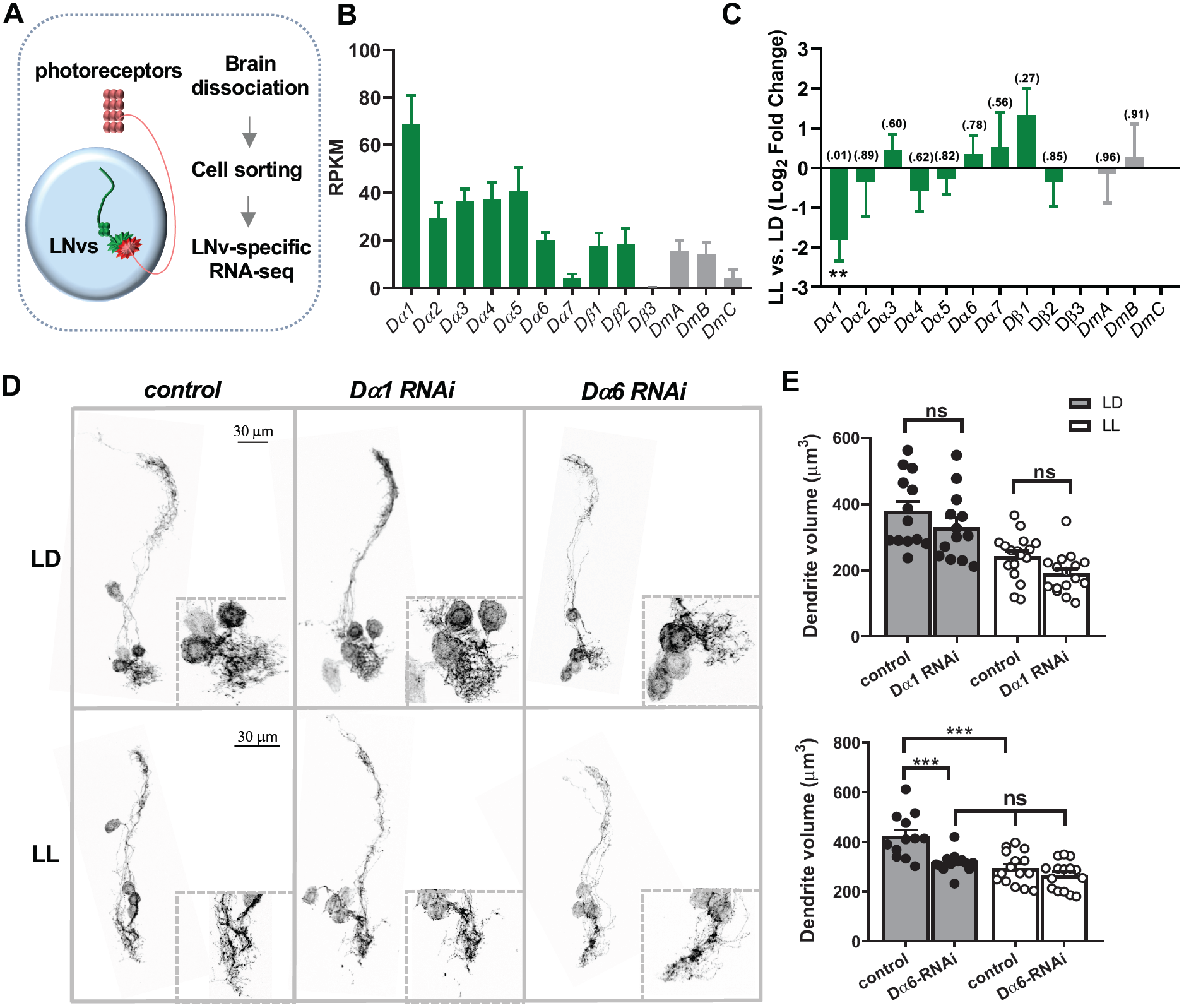
The nAchR subunits *Dα1* and *Dα6* are identified as candidate genes regulating larval LNv development. (**A**) A schematic diagram illustrating the LNv-specific RNA-seq analyses. LNvs receive synaptic inputs from larval photoreceptors. RNA-seq analyses were performed using LNvs sorted from 3^rd^ instar larval brains. (**B-C**) RNA-seq analyses reveal the relative expression levels of nicotinic (Dα1-7, Dβ1-3) and muscarinic AchR (DmA-C) in the larval LNvs and activity-induced changes. (**B**) Relative expression levels of AchR subunits in LNvs from larvae raised in the 12-hour light:dark (LD) condition. RPKM: Reads per kilobase per million. (**C**) Constant light (LL) conditions induce significant changes in the transcript level of *Dα1* without affecting other subunits. p-values are denoted above the bar. (**D**) Transgenic RNAi screens targeting nAchR subunits identified *Dα6* as a candidate gene affecting LNv dendrite development. Representative projected confocal images of mCD8::GFP-labeled LNvs and zoomed-in images of the dendritic region (dashed squares) are shown. Compared to controls, knocking-down *Dα6*, but not *Dα1* subunit, reduces the volume of LNv dendrites significantly in the LD condition (top panels) and eliminates the LL-induced decrease in LNv dendrite volume (bottom panels). (**E**) Quantification of the LNv dendrite volume. The light conditions and genotypes are as indicated. Sample size n represents the number of larvae tested. n=12-17. Error bars represent mean ± SEM. Statistical significance is assessed by one-way ANOVA with Tukey’s post hoc test. ns: not significant, ***: *p*< 0.001.

We also used RNA-seq analyses to test the expression of nAchR subunits under different light: dark conditions and to detect transcripts that undergo activity-dependent changes. Notably, Dα1 is the only subunit whose expression is significantly modified by chronically elevated levels of input activity: the exposure to constant light (LL) induced a roughly 3.5x reduction in expression compared to regular light: dark conditions (LD) (Figure 1C). Together, these results suggest that Dα1 is one of the main nAchR α subunits in LNvs and that its transcription level is regulated by activity.

To identify specific nAchR subunits involved in regulating LNv dendrite morphology or plasticity, we performed genetic screens using RNAi lines targeting all ten nAchR subunits. 3D visualization and quantification of the LNv dendrites showed that the LNv-specific RNAi knock-down of Dα6 induced a signification volume reduction and the loss of LL-induced dendrite plasticity (Figure 1D, E). Knocking down all other subunits, including Dα1, did not generate a consistent dendrite morphology phenotype (Figure 1D, E).

### Dα1 and Dα6 subunits are expressed in the larval LNvs

The combined results from the LNv-specific transcriptome analysis and transgenic RNAi screen suggested Dα1 and Dα6 as top candidates for follow-up studies. To confirm these two subunits’ expression in LNvs, we obtained enhancer-trap lines generated by the Trojan-Gal4 technique. This genetic approach introduces an artificial splice acceptor site at the 5’ end of a MiMIC insertion to terminate endogenous transcription while driving Gal4 expression (Diao et al., 2015). Enhancer trap lines *Dα1*^*MI00453-TG4.0*^ (Dα1-TG4) and *Dα6*^*MI01466-TG4.1*^ (Dα6-TG4) were then used to drive the expression of a membrane-targeted GFP, mCD8::GFP, and a nuclear marker, RedStinger (Nagarkar-Jaiswal et al., 2015a; Nagarkar-Jaiswal et al., 2015b). For both enhancer trap lines, we observed expression of the two markers in LNvs, indicating that the *Dα1* and *Dα6* loci are transcriptionally active in these cells (Figure 2). Notably, the intensity of Dα6-TG4*-*driven RedStinger was weaker than that of Dα1-TG4, in agreement with the relative expression levels reported by the RNA-seq study (Figures 2).

**Figure 2.**
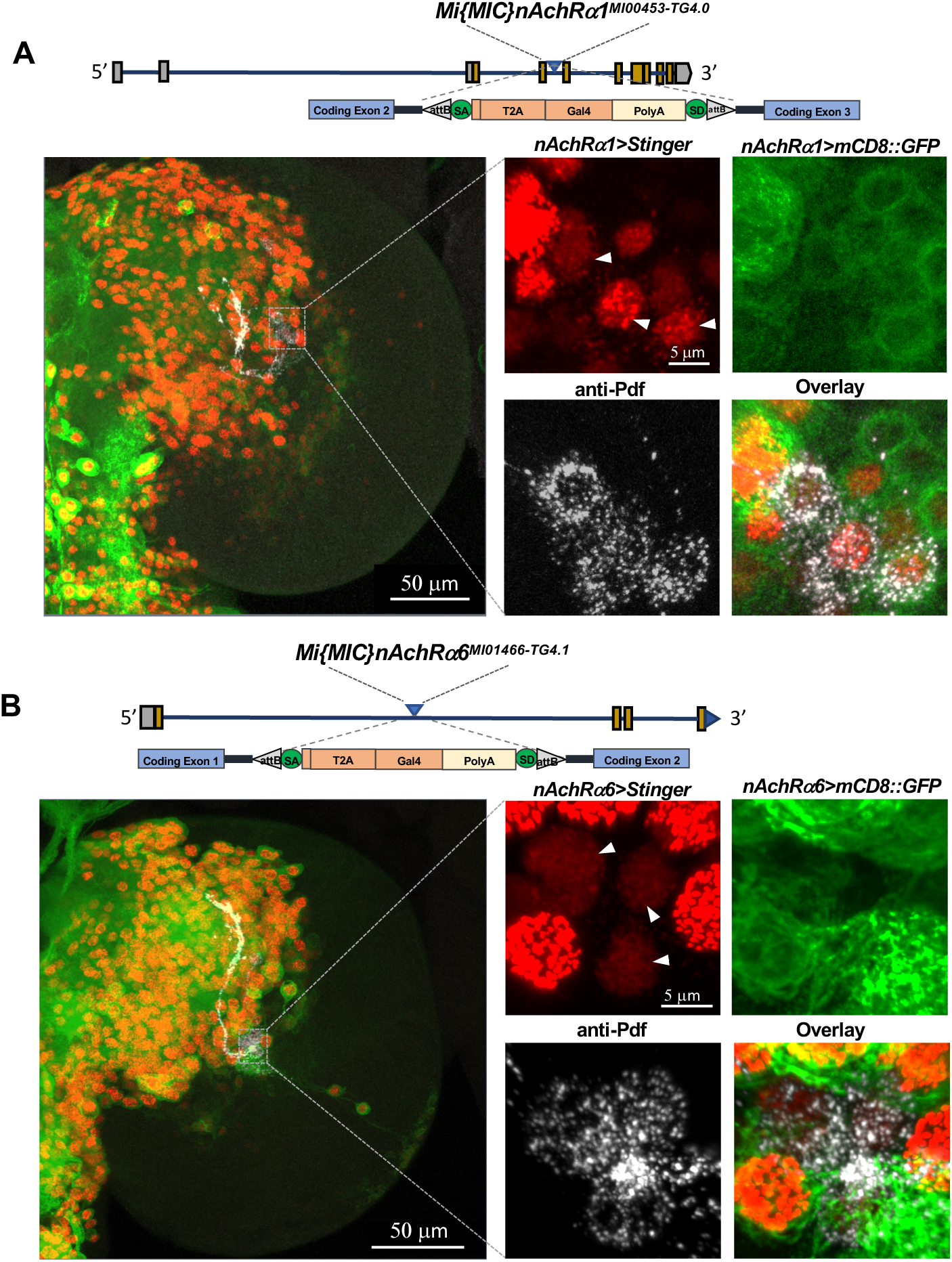
*Dα1* and *Dα6* subunits are expressed in the larval brain and LNvs. (**A-B**) The expression of *Dα1* and *Dα6* in the larval LNvs is confirmed using Gal4 enhancer trap lines. Top: Schematics of the Trojan-Gal4 constructs and the insertion sites within the *Dα*1 and *Dα6* genomic loci. Bottom: Representative projected confocal images displaying the broad distribution pattern of *Dα*1 and *Dα6* in the larval brain lobe (left) and in the LNvs (zoomed-in images, right). The Trojan-Gal4-driven expression of nuclear marker Stinger (red) and the membrane marker mCD8::GFP (green) are shown. LNvs are stained using anti-PDF antibody (white, arrow heads). Genotypes and scale bars are as indicated.

### *Dα6* is cell-autonomously required for LNv dendrite morphogenesis

Given the abundance of the Dα1 subunit in LNvs, it was surprising to see that Dα1 knock-down did not induce dendrite defects. To eliminate the possibility that our knock down was inefficient, we tested trans-heterozygous mutants consisting of two MiMIC alleles inserted in either a coding intron (*Dα1*^*MI00453*^) or an exon (*Dα1*^*MI11851*^) (Supplementary Figure 2A). Consistent with the knock-down experiments, we observed no changes in LNv dendrite volume in the Dα1 mutant under LD and LL conditions (Supplementary Figure 2B, C).

The contribution of Dα6 to LNv morphogenesis was also demonstrated by a mutant analysis using the null allele *Dα6*^*DAS1*^, which contains a G→A substitution that eliminates the first splicing event and produces a truncated polypeptide (Watson et al., 2010). As in the knockdown approach, Dα6 deficiency led to a significant reduction in the LNv dendrite volume and a loss of LL-induced dendrite plasticity (Figure 3A, B). Our RNAi knockdown experiments suggested that Dα6 regulates LNv dendrite development in a cell-autonomous manner (Figure 1D, E). To confirm this, we performed rescue experiments in the mutant background using the LNv-specific expression of a Dα6 transgene, which rescued the dendrite plasticity phenotypes and lessened the dendrite volume reduction observed in the *Dα6*^*DAS1*^ mutants (Figure 3A, B). Notably, the rescue transgene did not fully restore the LNv dendrite volume, which remained significantly lower than the volume of the control group, suggesting that *Da6* expressed outside LNv also plays a role in its development. Taken together, these morphological results suggest that *Da6* is required, in part *via* a cell-autonomous contribution, for LNv dendrite development.

**Figure 3.**
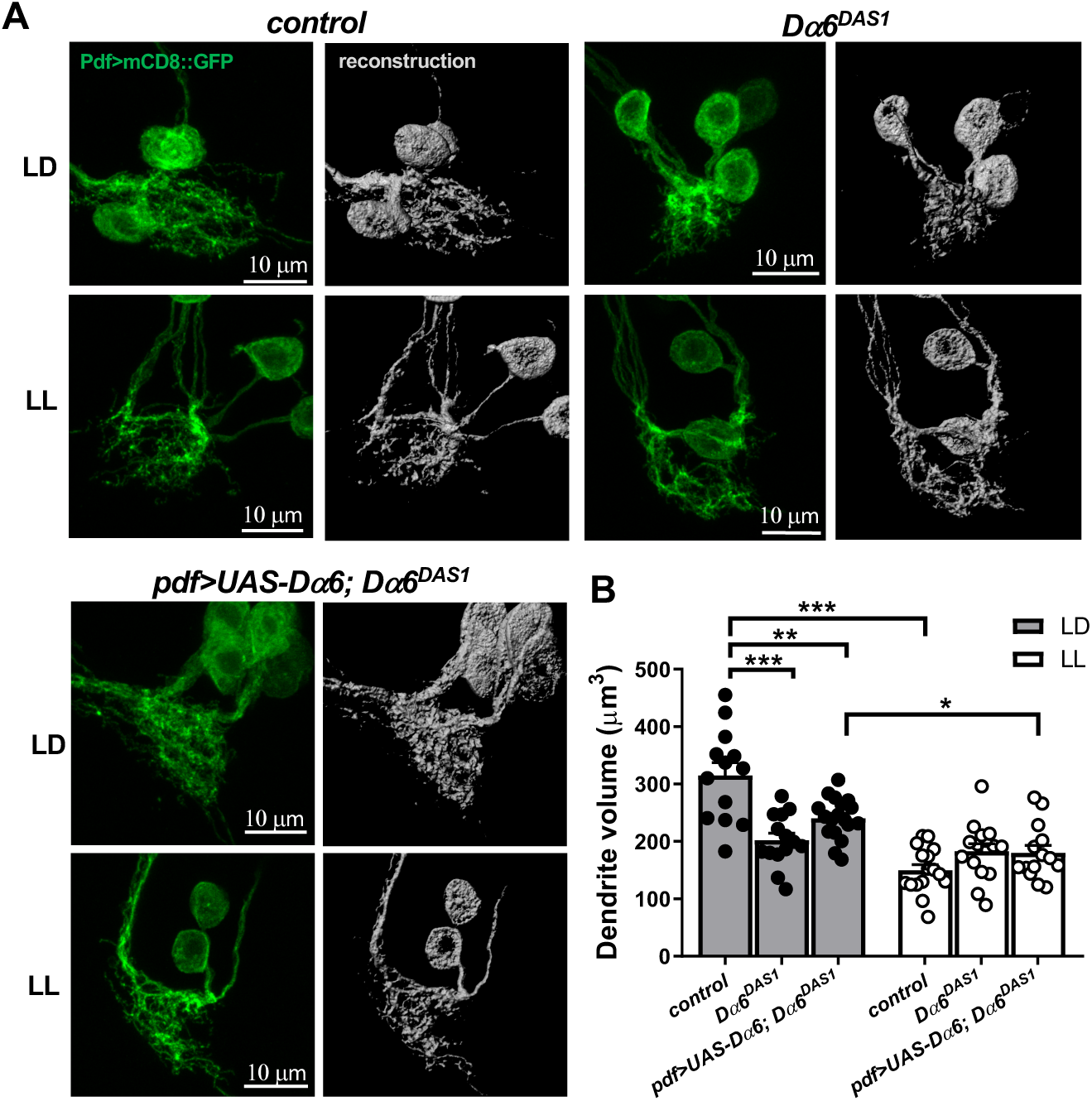
The *Dα6* subunit is required cell-autonomously for the structural development of LNv dendrites. (**A-B**) LNv dendrite development is affected in *Dα6* loss-of-function mutants (*Dα6*^*DAS1*^). Compared to controls, *Dα6*^*DAS1*^ mutants show a significant reduction in the LNv dendrite volume in the LD condition and a loss of LL-induced decrease in LNv dendrite volume. Both phenotypes are partially rescued by the LNv-specific expression of the *Dα6* transgene. (**A**) Representative projected confocal images of LNv dendrites and somas labeled by mCD8::GFP (left, green) and their 3D reconstructions (right, grey) are shown. The light conditions and genotypes are as indicated. (**B**) Quantification of the LNv dendrite volume. Sample size n represents the number of larvae tested. n=13-18. Statistical significance is assessed by one-way ANOVA with Tukey’s post hoc test. *: *p*< 0.05, **: *p*< 0.01, ***: *p*< 0.001. Error bars represent mean ± SEM.

### Both *Dα1* and *Dα6* subunits are required for synaptic transmission in LNvs

Although it was now clear *Dα6* played an important role in LNv dendrite morphogenesis, the functions of *Dα1* remained unknown. Thus, we examined these two nAchR subunits using physiological studies. LNvs are visual projection neurons that receive synaptic inputs from the larval photoreceptors and thus can be activated when the animal is subjected to light stimuli. Using genetically-encoded calcium indicator GCaMP6s and larval eye-brain explant preparations (Yuan et al., 2011), we performed two photon calcium imaging experiments on the LNvs while delivering 100ms light pulses with a 560nm laser. Light-evoked physiological responses were quantified by measuring changes in GCaMP6s signals recorded at LNvs’ axonal terminal region (Figure 4A, B).

**Figure 4.**
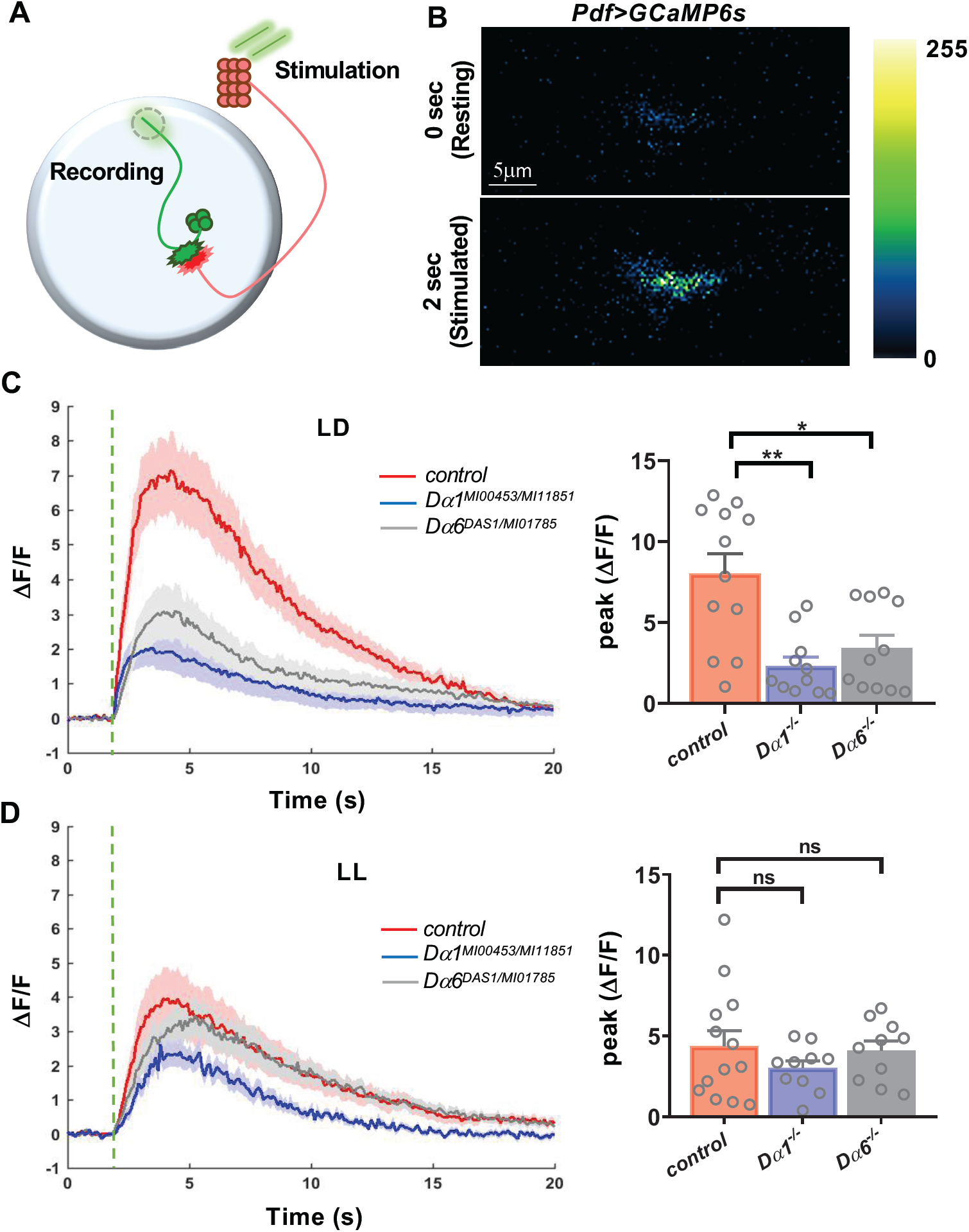
Both *Dα1* and *Dα6* subunits are required for light-evoked physiological responses in LNvs. (**A**) The schematic diagram illustrating the calcium imaging experiment in LNvs. Photoreceptor-mediated synaptic transmission is received by LNvs and measured in the axonal terminal region of LNvs (dashed circle). (**B**) Representative pseudo colored frames of *Pdf>GCaMP6s* recording demonstrating light-evoked increases of the GCaMP6s signals in the LNvs. (**C-D**) Light-elicited calcium responses in LNvs are severely affected in *Dα1* and *Dα6* loss-of-function mutants under the LD (**C**) but not the LL conditions (**D**). Genotypes of the trans-heterozygous mutants of *Dα1* and *Dα6* are as indicated. Left: Average calcium transients elicited by light pulses (beginning at green dashed lines). The shaded area represents SEM. Right: The quantifications of peak value of the changes in GCaMP signal induced by light stimulations (δF/F). Sample size n represents the number of larvae tested. n=10-13. Statistical significance is assessed by one-way ANOVA followed by Tukey post-hoc test. ns: not significant, *: *p*<0.05, **: *p*<0.01. Error bars represent mean ± SEM.

The first set of experiments was conducted in the LD condition using the *Dα1*^*MI11851*^*/Dα1*^*MI00453*^ and *Dα6*^*DAS1*^*/Dα6*^*MI01785*^ trans-heterozygous loss-of-function mutants. Remarkably, the light-elicited calcium response in both *Dα1* and *Dα6* mutants was significantly reduced compared to the control flies. Response amplitudes, represented by the peak level of the ΔF/F, were only about 30% of control for the *Dα1* mutant and 40% of control for the *Dα6* mutant, although there was no statistical difference between these two groups (Figure 4C).

Previously, it has been shown that LNvs of LL-cultured larvae undergo not only a reduction in dendrite volume, but also a dampened light-evoked calcium response (Yuan et al., 2011). Interestingly, both *Dα1* and *Dα6* mutants cultured in LL display a similar reduction in physiological response as the wildtype larvae (Figure 4D). This inability for LL condition to enhance the mutants’ phenotype suggests that the environmental influence introduced by LL and the genetic deficits generated by Dα1 and Dα6 mutations share the same cellular target, possibly the number and/or strength or the synapses.

While the loss of *Dα6* likely affects synaptic transmission by regulating LNv postsynaptic development, the strong impact generated by *Dα1* deficiency on the LNvs’ physiological properties is striking given the lack of phenotype in dendrite morphology. These observations suggest excitatory synaptic transmission and dendrite development are separately regulated, resembling a finding made in the glutamatergic synapses in the vertebrate hippocampus, where eliminating synaptic transmission produced no obvious perturbations of neuronal morphology (Lu et al., 2013).

### *Dα1* and *Dα6* subunits are transcriptionally regulated during larval development

Our morphological and physiological evaluations of the two nAchR subunits revealed their distinct functions: Dα6 is important for the structural development of the postsynaptic compartment and supports synapse formation and dendrite growth, whereas Dα1 mainly contributes to synaptic transmission. These functional differences between nAchR subunits resemble the ones observed in vertebrate NMDA and AMPA glutamatergic receptors. In the vertebrate system, the maturation of excitatory glutamatergic synapses is regulated by the ratio of AMPA/NMDA receptors (Wu et al., 1996). To test whether similar developmental regulation of receptor subtypes is also present in the *Drosophila* central cholinergic synapse, we sought to examine the levels of two nAchR subunits in young vs. mature LNvs.

To analyze the transcript level of nAchR subunits in the LNvs, we performed qFISH experiments on acutely dissociated LNvs using subunit-specific probes (Yin et al., 2018) (Figure 5A). This set of experiments confirmed the expression of both subunits in the LNvs and indicated that the expression level of Dα1 is low in young LNvs, between 48 and 72 hours after egg laying (48-72 hr AEL) but is significantly upregulated later in development (120 hr AEL) (Figure 5A). In contrast, the level of Dα6 in LNvs declined significantly during larval development (Figure 5B). Notably, we also observed that the excessive synaptic input generated in the LL condition had a strong effect: the developmental upregulation of Dα1 was eliminated, removing the significant difference observed between the two developmental stages in the normal LD condition.

**Figure 5.**
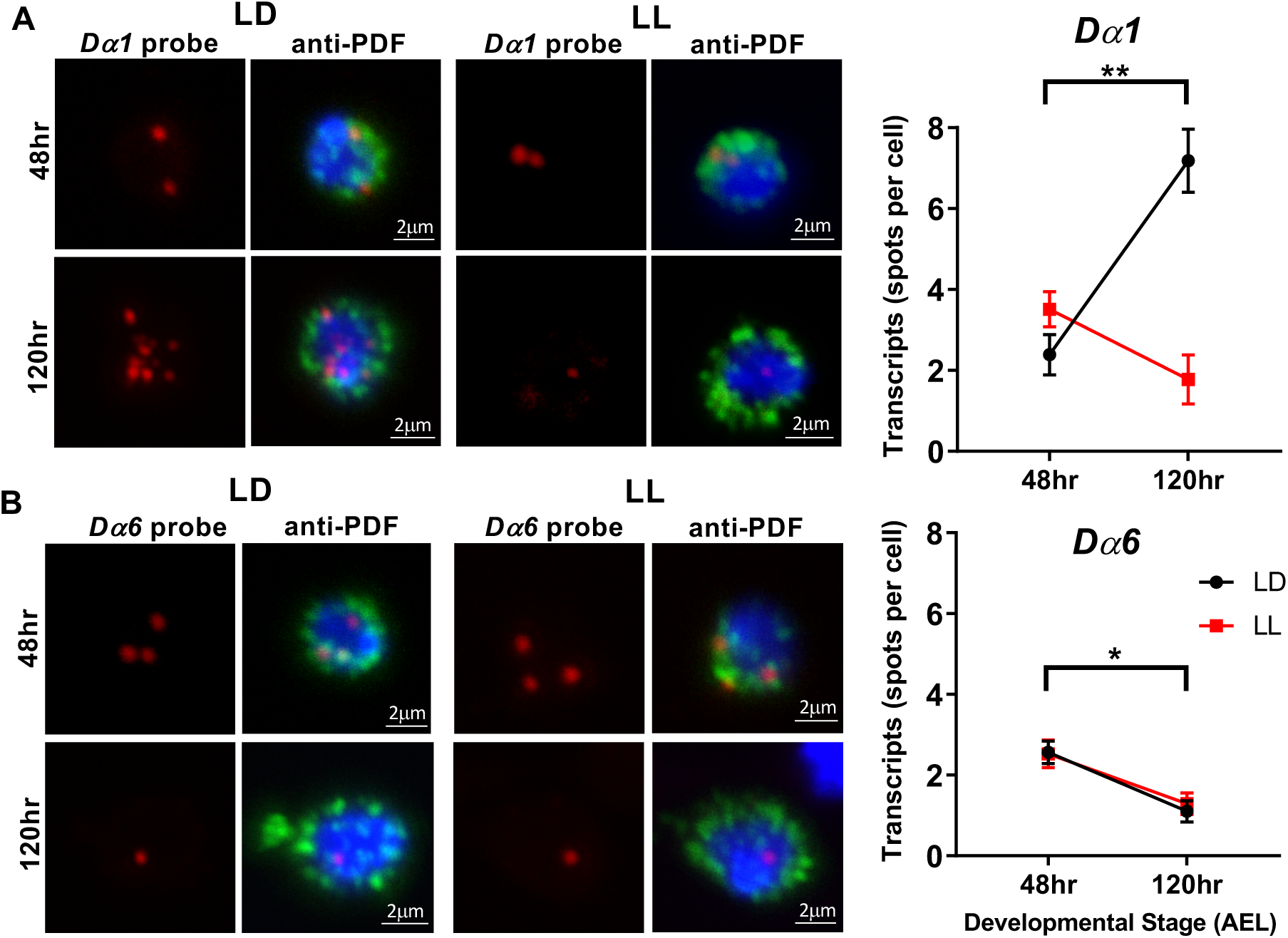
*Dα1* and *Dα6* subunits are transcriptionally regulated during larval development. (**A**) The transcript level of *Dα1* in LNvs is regulated during development and influenced by activity. LL conditions eliminate the upregulation of *Dα1* transcripts at 120hr after egg laying (AEL). Left: Representative projected confocal images of qFISH experiments labeling *Dα1* mRNA transcripts (red) in acutely dissociated LNvs (green) are shown. The LNvs were collected from larvae at two different developmental stages. Cell nuclei are stained with DAPI (blue). Right: Quantifications of the *Dα1* transcript level revealed a significant increase at the later developmental stage when larvae are cultured in the LD condition (black) but not the LL condition (red). n represents number of dissociated LNvs. n=18-62 in all groups. (**B**) *Dα6* transcript level in LNvs is downregulated during larval development and is not sensitive to input activity. Quantifications revealed a significant reduction of LNv transcripts in both LD and LL conditions. n represents number of dissociated LNvs. n=39-54 in all groups. Statistical significance is assessed by one-way ANOVA followed by Tukey post-hoc test.. *: *p*<0.05, **: *p*<0.01. Error bars represent mean ± SEM. Scale bars are as indicated.

The qFISH experiments revealed that Dα1 and Dα6 had different temporal profiles of expression in LNvs, and different sensitivity to chronic alterations in activity. These observations are consistent with results obtained from the RNA-seq analyses (Figure 1B, C) and support the notion that Dα1 and Dα6 have different functional roles in the structural and functional development of LNvs and activity-induced plasticity.

### *Dα6* is essential for cholinergic synapse development in larval LNvs

Based on the results obtained thus far, we propose that Dα1 and Dα6 are preferentially expressed at distinct time periods to meet the changing needs of developing and mature LNvs. Our previous studies indicate that immature LNvs, from the 2^nd^ to early 3^rd^ instar stages and between 48-72hr AEL are characterized by highly dynamic dendritic filopodia that support synapse formation. This developmental process reaches its peak at 72hr AEL and then declines drastically by the mid-3^rd^ instar stage (Sheng et al., 2018). Here, our data indicate that the end of the synaptogenesis period temporally correlates with the developmental reduction of *Dα6* transcripts. This raises the possibility that *Dα6* is the receptor subunit responsible for establishing the cholinergic synapse on the dendrites of young LNvs, while *Dα1* mainly functions in mature LNvs and supports synaptic transmission.

To test this hypothesis, we examined putative synaptic contacts originating from the Bolwig’s nerve (BN), which contains axonal projections of the larval photoreceptors, onto the LNv dendrites, using an mCherry-tagged presynaptic active zone marker, Bruchpilot (Brp), driven by the photoreceptor-specific enhancer (Rh5,6-Brp::mCherry) (Mosca and Luo, 2014; Sheng et al., 2018). In addition, using 3D reconstructions of mCD8::GFP-labeled LNv dendritic arbors and the Rh5,6-Brp::mCherry labeled presynaptic terminals, we quantified the LNv dendrite volume, the total number of presynaptic terminals generated by the BN, and the number of the Brp puncta directly contacting LNv dendrites (Figure 6A-C) (Sheng et al., 2018).

**Figure 6.**
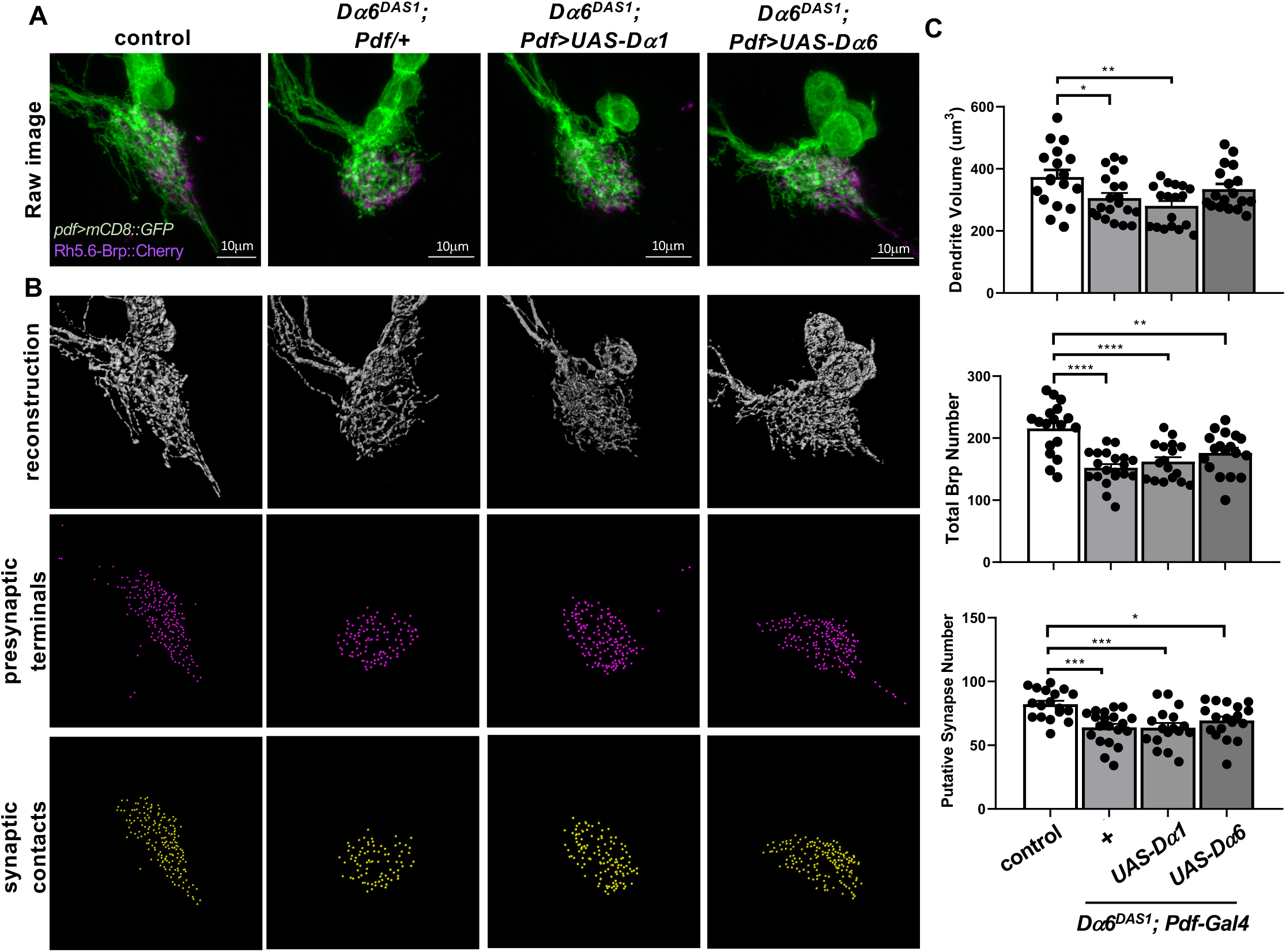
*Dα1* and *Dα6* have non-redundant functions in LNv dendrite and synapse development. (**A-C**) *Dα6* deficiency leads to reduced LNv dendrite volume, total number of presynaptic terminals and synaptic contacts between LNv and presynaptic photoreceptors. LNv-specific expression of a *Dα6* transgene partially rescued the dendrite volume phenotype in the *Dα6*^*DAS1*^ mutants, while the expression of a *Dα1* transgene had no significant effect. (**A**) Representative projected confocal images of the LNv dendrites (green) co-labeled with *Rh5,6-Brp:mCherry* (magenta). (**B**) The LNv dendrite volume (gray), the number of BO presynaptic terminals in the LON region (*Rh5,6-Brp* puncta, magenta spots) and the number of Brp punctae contacting the LNv dendrites (synaptic contacts, yellow spots) were analyzed using 3D reconstructions. Representative images for samples with the indicated genotypes are shown. (**C**) Quantification of the volume of LNv dendrites (Top), the total number of *Rh5,6-Brp* puncta (Middle) and the number of putative synaptic contacts (Bottom) revealed significant deficiencies in *Dα6*^*DAS1*^ mutants and the partial rescue generated by LNv-specific expression of a *Dα6* transgene. n=12-17 in all groups. Statistical significance was assessed by one-way ANOVA followed by Tukey’s *post hoc* test. *: *p*<0.05, **: *p*<0.01. ***: *p*<0.001 Error bars represent mean ± SEM. Scale bars and genotypes are as indicated.

Consistent with our previous observations, *Dα6*^*DAS1*^ mutants showed a significant reduction in dendrite volume, as compared to the controls. In addition, the LNv-specific expression of a *Dα6* transgene partially rescued this phenotype (Figure 6A-C), while expressing a *Dα1* transgene in a similar fashion did not modify the *Dα6* mutant phenotype. Together, these results suggest that these two receptor subunits have distinct molecular properties that support separate functions in regulating LNv development.

The quantifications of Rh5,6-Brp::mCherry puncta also revealed that the D*α6* deficiency had a strong effect on the overall number of presynaptic terminals made by the BN in the larval optic neuropil (LON) (Figure 6A-C), which includes the putative synaptic contacts between the BN and LNvs as well as other postsynaptic target neurons downstream of the larval photoreceptors (Sheng et al., 2018; Sprecher et al., 2011). In conjunction with the broad expression of D*α6* we observed in the larval CNS (Figure 2B), and a previous report on its role in regulating the nAchR-specific synapse protein Hikaru Genki (Hig) in the adult mushroom body calyx (Nakayama et al., 2016), this result suggests that D*α6* is also required for synaptogenesis in other types of neurons in the larval CNS. It was not surprising, therefore, that the rescue experiments using LNv-specific expression of either the Dα6 or Dα1 transgene did not produce a significant change in the total number of Brp punctae in the Dα6 mutant background (Figure 6A-C).

Lastly, we examined putative synaptic contacts between the BN and LNv by determining the number of Brp puncta that are in direct contact with LNv dendrites. This result showed a trend similar to that of the total Brp punctae in the *Dα6* mutant and rescue groups (Figure 6A-C), suggesting that the number of synapses formed on the LNv dendrite is regulated by the *Dα6* subunit through both autonomous and non-autonomous contributions.

Taken together, our results indicate that, besides regulating LNv dendrite development, *Dα6* is also required for establishing the synapse between the larval photoreceptor and LNv dendrites. This function likely relies on *Dα6* expression in both LNvs and non-LNv neurons. In addition, the inability of a temporally mis-expressed *Dα1* transgene to substitute for the loss of *Dα6* in rescue experiment supports the functional distinction between the two subunits. Currently, our understanding of the biophysical roles of insect nAchR subunits is lacking relative to their vertebrate counterparts. However, the amino acid sequence alignments revealed clear differences between Dα1 and Dα6 in their key functional motifs, including the Cys-loop that is critical for ligand binding-induced channel opening and the TM3-TM4 intracellular loop, which is a common site on nAchRs for post-translational modifications and protein-protein interactions (Jones et al., 2010) (Supplementary Figure 3).

## Discussion

Our genetic studies analyzed the role of nAchR signaling in regulating central cholinergic synapse development and dendrite morphogenesis. The results indicate that larval LNvs rely on specific nAchR subunits for structural and functional maturation of the postsynaptic apparatus. The distinct functional roles of the Dα1 and Dα6 subunits are supported by transcriptional programs regulating their expression and alter nAchR receptor composition in young vs. mature LNvs. While Dα6 is required for synapse development and is downregulated after the synaptogenesis period, Dα1 is necessary for mediating synaptic transmission and is upregulated in mature LNvs. Together, our data support transcriptional controls of functionally distinct nAchR subunits as a critical element regulating the development of central cholinergic synapses.

### *Dα6* as the molecular candidate for nAchR-mediated calcium-dependent synaptic plasticity

Unlike vertebrate nAchR genes whose physiological and molecular properties have been studied sufficiently to allow a detailed description of the conductance and kinetics of each receptor subtype, insect nAchR genes are not well understood. In the vertebrate, Chrna7 homo-pentamers are highly permeable to Ca^2+^ ions, more so even than glutamatergic NMDA receptors, and display a rapid desensitization following ligand-gated activation. Conversely, α4β2, α2β4 and other hetero-pentamers have a comparatively low Ca/Na conductance and desensitize slowly (Albuquerque et al., 2009; Bertrand et al., 1993; Fucile, 2004; Seguela et al., 1993). That these subunits show differential conductance of cation species is important because, in contrast to the fast, depolarizing effect of Na^+^ ions, Ca^2+^ ion flux is crucial to the induction of a host of calcium-dependent intracellular signaling pathways that could lead to long-term changes in dendrite development and postsynaptic specification, most notably, the clustering of “late-stage receptors” (Cooke and Bliss, 2006). Based on our analyses, we propose that Dα6 mediates the activation of calcium-dependent signaling pathways that guide the structural development and activity-induced plasticity in the LNv dendritic arbor.

In insects, the involvement of calcium-permeable, excitatory neurotransmitter receptors in neuronal plasticity has been well demonstrated. In both honeybee and *Drosophila*, nAchRs have been implicated in long-term memory and learning (Campusano et al., 2007; Gauthier et al., 2006). In addition, previous studies showed that the activation of nAchRs induces calcium-dependent plasticity in Kenyon cells (Campusano et al., 2007). *Da6*, which is expressed early in development in immature LNvs, and which shares close homology to the Ca^2+^-conducting mammalian Chrna7, is a suitable molecular candidate for coordinating calcium-dependent changes that regulate synapse development and long-term plasticity. These functions, however, may be performed in a redundant fashion by Dα5 and Dα7, both of which are also homologues of vertebrate α7. To elucidate the cellular and molecular mechanisms underlying nAchR-mediated calcium-dependent plasticity in the *Drosophila* CNS, future studies will be centered on resolving the physiological properties and tissue distributions of these three subunits, deficits generated by different combinations of their mutants, and potential *in vitro* characterizations of the conductance and kinetics of these subunits.

### Subunit-specific regulation of *Dα1* and *Dα6*

Our current model suggests that the developmental switching of nAchR subunits is a signature event accompanying the maturation of central cholinergic synapses. However, it is unclear whether the changes in nAchR composition are instructive or permissive for the maturation of postsynaptic specifications in the LNvs. To address this question, a deeper molecular understanding of subunit-specific regulatory mechanisms is needed.

Previous studies in both vertebrate and *Drosophila* primary neuronal cultures have demonstrated that the level of mammalian nAchR α7 or Dα7 can be modulated by neuronal activity, possibly through post-transcriptional mechanisms (Brumwell et al., 2002; Ping and Tsunoda, 2011). Our studies, however, including the initial RNA-seq analyses and the subsequent qFISH experiments, indicate that the levels of Dα1 and Dα6 subunits are regulated by distinct transcriptional programs. While expression of *Dα1* is modulated by visual experience: excessive light input suppresses the transcriptional upregulation that normally occurs through the course of larval development, the *Dα6* transcript is downregulated in mature LNvs regardless of changes in light conditions. Future studies identifying the distinct transcriptional programs controlling the differential expression of *Dα1* and *Dα6* will facilitate our understanding of regulatory mechanisms underlying synapse and dendrite development, as well as the impact of sensory experience on the transcriptional programs organizing activity-induced plasticity events (Chen et al., 2016; Flavell and Greenberg, 2008; Malik et al., 2014; Spiegel et al., 2014).

Our studies only examined the transcript levels of the Dα1 and Dα6 subunits, and therefore does not address the possibility that concurrent post-transcriptional regulatory mechanisms are also involved in the processing, maturation and synaptic integration of nAchR subunits during cholinergic synapse development. A number of nAchR-associated accessory proteins, such as RIC-3, 14-3-3 proteins, LRP4 and Nacho, have been identified in vertebrate, *C. elegans* and *Drosophila* systems (Gu et al., 2016; Jones et al., 2010; Mosca and Luo, 2014; Sadahiro et al., 2016). These auxiliary proteins are required for the proper folding and trafficking of nAchR subunits and could act as additional regulatory elements to ensure the preferential distribution of *Dα1* and *Dα6* subunits at their respective synaptic sites.

### Developmental switching of the neurotransmitter receptor subtype as a common feature for synapse maturation

During larval development, the expression level of Dα1 in LNvs is upregulated while the expression level of Dα6 is downregulated, suggesting a change in nAchR receptor composition during LNv maturation. The distinct functional roles and temporal regulation of the two nAchR subunits resemble developmental switches described for both glutamate and glycine receptors in the developing vertebrate CNS (Ben-Ari et al., 1997; Wu et al., 1996).

In the vertebrate system, the maturation of excitatory glutamatergic synapses is regulated by the shifting ratio of AMPA/NMDA receptors (Pratt et al., 2016). Electrophysiological recordings from *Xenopus* tectal neurons indicate that neurotransmission in immature neurons is predominantly mediated by NMDA receptors. As neurons mature, the CaMKII-dependent trafficking of AMPA receptors leads to an increased AMPA/NMDA ratio, which enhances neurotransmission and synapse strength, contributing to the physiological maturation of the synapse (Pratt et al., 2016; Sin et al., 2002; Wu et al., 1996). Additionally, the molecular mechanism responsible for long-term potentiation (LTP) is also known to involve a change in receptor type. In this case, synaptic connections, such as those found in the hippocampus, become strengthened through NMDA-dependent recruitment of AMPA receptors, which relies on intracellular calcium influx (Cooke and Bliss, 2006; Sheng and Kim, 2011; Volianskis et al., 2015).

Remarkably, similar molecular changes were also observed during the maturation of the postsynaptic apparatus of the cholinergic vertebrate NMJ, where the embryonic form of nAchR containing a gamma subunit (α2βγδ) is replaced by nAchRs containing an epsilon subunit (α2βεδ) *via* transcriptional regulation (Sanes and Lichtman, 2001). In combination with these previous findings, our study in a central cholinergic synapse of the *Drosophila* system supports the developmental switching of neurotransmitter receptor composition as a common mechanism underlying the maturation and activity-dependent regulation of synapse development and function.

## Methods

### Fly culturing

All flies were maintained in standard cornmeal medium supplemented with yeast paste and in controlled incubators set at 25°C and 60% humidity. Animals subjected to the LD condition were reared at alternating 12hour light-12hour dark cycles and those subjected to the LL condition receive 24hours of light. All larvae used for experiments were taken at the wandering 3^rd^ instar stage, with the exception of early stage larvae in Figure 5 where animals were dissected between 48hr to 72hr after egg laying.

### *Drosophila* stocks and genetics crosses

The genetics of knockdown experiments used in Figure 1, including the preliminary screen, were performed with *Pdf-Gal4, UAS-mCD8::GFP; UAS-Dicer* (Bloomington 6899, 5137 and 24650) and *Dα1* and *Dα6* RNAi stocks (Bloomington 28688 and 25835). The Trojan-Gal4 studies in Figures 2 used *yw;+;Dα1*^*MI00453-TG4.0*^*/TM6C* and *yw; Dα6*^*MI01466-TG4.1*^;+ (Bloomington 66780 and 76137). The mutant study in Figure 3 was performed with the *Dα6*^*DAS1*^ (Bloomington 9685) and *UAS-Dα6* transgenic flies made in this study. The mutant study in Supplementary Figure 3 used *Pdf>mCD8::GFP* and the MiMIC lines *yw;Dα1*^*MI11851*^ and *yw;;Dα1*^*MI00453*^ (Bloomington 56462 and 42295,). The physiological studies of Figure 4 used the stocks *Pdf-LexA* (Yuan et al., 2011) driving *LexAop-GCaMP6s* (Bloomington 44590) as well as an additional MiMIC mutant line *yw; Dα6*^*MI01785*^;+ (Bloomington 37925). Quantitative FISH analyses of Figure 5 used larvae with the genotype *Pdf-Gal4, UAS-mCD8::GFP*. The experiment shown in Figure 6 was performed using *Pdf-Gal4, mCD8::GFP; Rh5.6-Brp::mCherry* expressed in the *Dα6*^*DAS1*^ mutant background with or without UAS-Dα1 and UAS-Dα6 transgenes.

### Generation of transgenic *Dα6*-overexpression rescue lines

The coding sequence of the *Drosophila Dα6-*RE, the isoform most highly expressed in the larval LNv determined by the RNA-seq analyses (Yin et al., 2018), was amplified using the forward primer 5’-ACAGATCTTGCGGCCGCATGGACTCCCCGCTGCCAGCGTCG-3’ and the reverse primer 5’ ACAAAGATCCTCTAGATTATTGCACGATTATGTGCGGAGCG 3’. The resulting fragment was digested with NotI and XbaI and cloned into the vector pUAST. After verifying the *Da6* sequence by sequencing and restriction digestion, plasmids were injected by Rainbow Transgenic Flies (Camarillo, CA) and transformed by random P-element insertion, followed by standard screening and balancing.

### Quantitative analysis of dendrite volumes and putative synaptic contacts

Larval brains were dissected in 1x PBS, fixed in 4% PFA (paraformaldehyde in 1x PBST) for 40 minutes at room temperature, washed three times with PBST (0.3% Triton X-100 in 1x PBS), equilibrated for 15 minutes and mounted in antifade medium (SlowFade™ Antifade Kit, Thermo Scientific S2828). LNvs were imaged with a Zeiss 700 confocal microscope with a 40x oil objective and Z-stacks were obtained by taking serial sections at ∼0.5μm thickness, with a typical x-y-z resolution of .09μm × .09μm x .49μm. For volumetric analysis, confocal Z-stacks of GFP-labeled dendrites were imported to the software Imaris and reconstructed using the surface module. All individual volumes generated using a predetermined threshold were then summed to create a final dendrite volume.

The Brp-mCherry punctae representing discrete presynaptic terminals were determined using the Spots module of Imaris to identify spherical volumes of 0.6μm in diameter. The intensity threshold, used in previous experiments, was determined by manual counting (Sheng et al., 2018). The “Spots close to surface” Imaris extension was subsequently used to determine all punctae whose center is within 0.3μm of the reconstructed LNv dendrite surface. This effectively reveals the number of BO-LNv synapses in this region. All experiments generating dendrite volume and/or synapse number data were blinded, by concealing the identity of each group, before the image processing step to avoid subjective bias during quantification.

### Immunohistochemistry

For the Trojan-Gal4 studies, brains dissected in 1x PBS were fixed and washed as above and then incubated with primary antibody (1:10 mouse anti-Pdf, DSHB, Iowa City, Iowa) in 5% normal donkey serum (NDS) in 1x PBST, at 4 °C overnight. Samples were washed three times with PBST and incubated in secondary antibody (1:500 Goat anti-mouse Alexa647^+^, Thermo Fisher, Waltham, Massachusetts) for 2 hours at room temperature. Brains were then mounted and imaged. For *in situ* staining protocol, procedure is identical with the following modifications: cells are incubated with DAPI for 30 seconds before first PBST wash and after final PBST wash; primary antibody incubation uses BSA (1% Bovine serum albumin in DPBS) in place of NDS; the secondary antibody used is Alexa488^+^ Goat anti-mouse (1:2000 Thermo Fisher, Waltham, Massachusetts).

### Calcium imaging analyses

The protocol used here is adapted from previous publications (Sheng., et al., 2018, Yuan., et al. 2011). Eye-brain explants, including the intact Bolwig’s Organ, optic tract, eye disks and brain lobes, were taken from wandering 3^rd^ instar larvae expressing Pdf-LexA driving LexAop-GCaMP6s. After dissection in 1x PBS, samples were directly mounted in an external saline buffer (120mM NaCl, 4mM MgCl2, 3mM KCl, 10mM NaHCO3, 10mM Glucose, 10mM Sucrose, 5mM TES, 10mM HEPES, 2mM Ca2+, PH 7.2) for live *ex-vivo* imaging. Imaging was performed using a Zeiss LSM 780 confocal microscope in combination with a Coherent Vision II two-photon laser tuned to 920nm for GCaMP excitation. Samples were recorded with a 40x water objective lens and 3x optic zoom. The frame rate was set to 100ms for a total of 1000 frames over a 100s recording session. GCaMP signals at the axon terminal of LNv were recorded with a 256×90 pixel resolution. Each recording session included two 100ms 561nm light stimulations separated by 40s to allow for complete measurement of calcium response and return to baseline fluorescence. Fluorescence change following each stimulation was calculated by ΔF/F_0_, where ΔF represents the change in fluorescence from baseline levels and F_0_ represents the average fluorescence from the 20 frames immediately preceding stimulation. Only one recording was used per sample and traces and values in figures are derived from the data obtained from the second stimulation but are comparable to that of the first stimulation.

### Quantitative Fluorescent *In Situ* Hybridization assay (qFISH)

The qFISH protocol is based on (Yin., et al., 2018) with slight modifications. Larvae were dissected in DPBS (DEPC treated 1x PBS) with the ventral nerve cord removed. The brain lobes were subsequently broken with hypodermic needles and digested in a mixture of 200μL 1x Collagenase/1x Dispase and 50μL 1x Liberase at 25°C for 20-30 minutes. Enzyme treated brains were titrated 100 times until no substantial fragments remained. Digestion reaction was terminated by addition of 1mL SM3-FBS (Schneider’s insect medium with a 100μg/mL Penicillin-Streptomycin-Kanamycin antibiotic cocktail and 5% Fetal Bovine Serum). Cells/tissue fragments were centrifuged at 950g for 5 minutes at 10°C. All but 50mL of supernatant was removed and 50mL new SM3-FBS was added, mixed and then incubated at RT in a chamber pre-treated with .25mg/mL Concanavalin-A for 15 minutes to allow for cell adherence, followed by fixation in 4% PFA/PBS. Chambers were washed with PBS three times and subjected to serial ethanol washes (50%, 70%, 100%, 70% and 50%), then washed with PBS and incubated in a 1:15 Protease III solution (in 1x DPBS). Chambers were then rinsed with PBS before undergoing the hybridization portion of the protocol, which is performed using the RNAScope kit (ACD Bio, Catalog no. 320850, 820851) supplied with *Dα1* or *Dα6* mRNA hybridization probes (Catalog no. 555611, 515521). Dissociated cells were incubated with either *Dα1* or *Dα6* probes at 40°C for 2 hours. After the washing and amplification steps following the manufacturer’s protocols, the dissociated cells were stained using the anti-PDF antibody and mounted with vector shield for imaging.

### Statistical analysis

For all quantitative analyses, more than 10 biological repeats from 2 to 3 technical replicates are included. The quantifications of dendrite volume and synaptic contacts were performed blindly. For calcium imaging experiments, the data analyses and graphing were performed using a custom written MATLAB script to reduce human biases. The raw values were imported to Graphpad Prism 8.0 and analyzed using one-way ANOVA followed by Tukey’s multiple comparisons post-hoc test, comparing all means against each other. For calcium imaging experiments, maximum ΔF/F for each sample was calculated and graphed by a custom written MATLAB script to represent the amplitude value. For qFISH experiments, spots were manually counted by surveying full z-stack images, and only isolated cells, in which the spots can be clearly distinguished, were included. All figures depict data in a bar plot representation with individual values, except for Figure 5 which hides individual values due to large sample sizes (N>18). Levels of significance were established at *: *p*<.05, **: *p*<0.01, ***: *p*<0.001. Error bars represent standard errors of the mean.

### Sequence alignment

The phylogenetic tree was generated by including all 16 known human nAchR subunit genes and all ten *Drosophila* nAchR subunit genes. For human orthologs, the amino acid sequence used in the alignment was chosen from the primary listed isoform on the UniProt database. For *Drosophila* orthologs, polypeptide sequences were taken from the FlyBase database. In the event that multiple coding isoforms exist or are predicted, the isoform which contributes the verified peptide species, or the predominant isoform expressed, was selected. Phylogenetic comparisons and tree reconstructions were made by the MABL online tool using “One click” mode.

### Data availability statement

High throughput sequencing data presented in Figures 1 were previously deposited with the accession number GEO: GSE106930 (Yin *et al*. 2018). All raw data and detailed statistical analysis information can be found in the Source data file.

## Acknowledgments

We thank Bloomington Stock Center and the Gene Disruption Project for providing transgenic and mutant fly lines; and Mark Stopfer and Benjamin White for helpful discussions and comments on manuscripts. This research is supported by the intramural research program of National Institutes of Health. Project number 1ZIANS003137.

## Author contributions

J.R. and Q.Y. designed the experiments. J.Y. performed RNA-seq and bioinformatics analyses. J.R. performed data collection and analyses with the help from J. Y., E.S., C.L., and C. S. J.R. and Q.Y. wrote the manuscript.

---

**Supplementary Figure 1.**
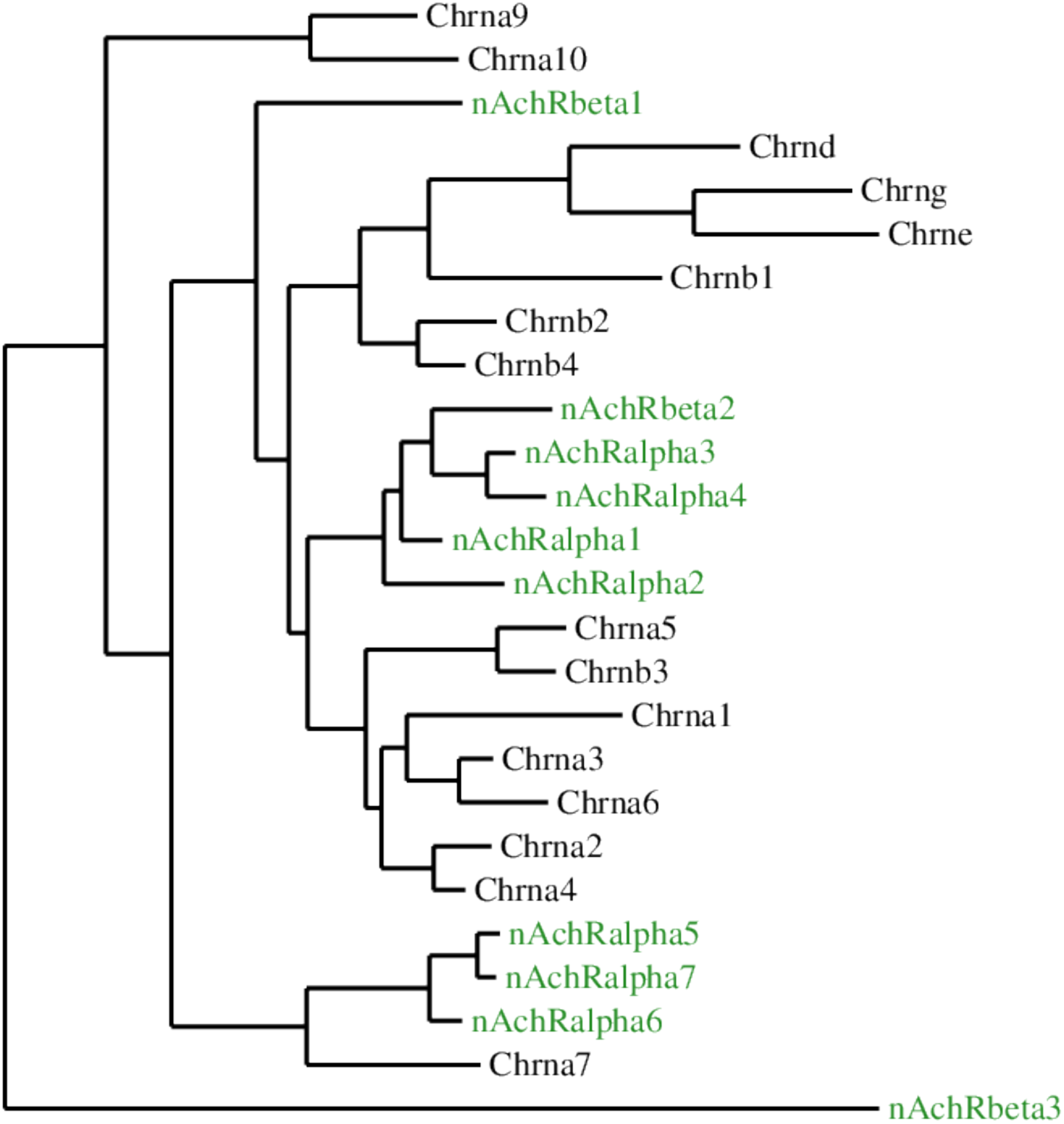
Alignment of the *Drosophila* nAchR gene family reveals distinct clusters. Phylogenetic tree depicting amino acid similarity between the ten *Drosophila* nAchR subunit genes (green) and the 16 human homologs (black) is shown. The fly nAchR gene family forms distinct clusters: Dα5-Dα7 cluster with human Chrna7 whereas Dβ1, Dβ2 and Dα1-Dα4 cluster with the remaining Chrna genes. Dβ3 is identified as the outgroup and is likely a species-specific subunit.

**Supplementary Figure 2.**
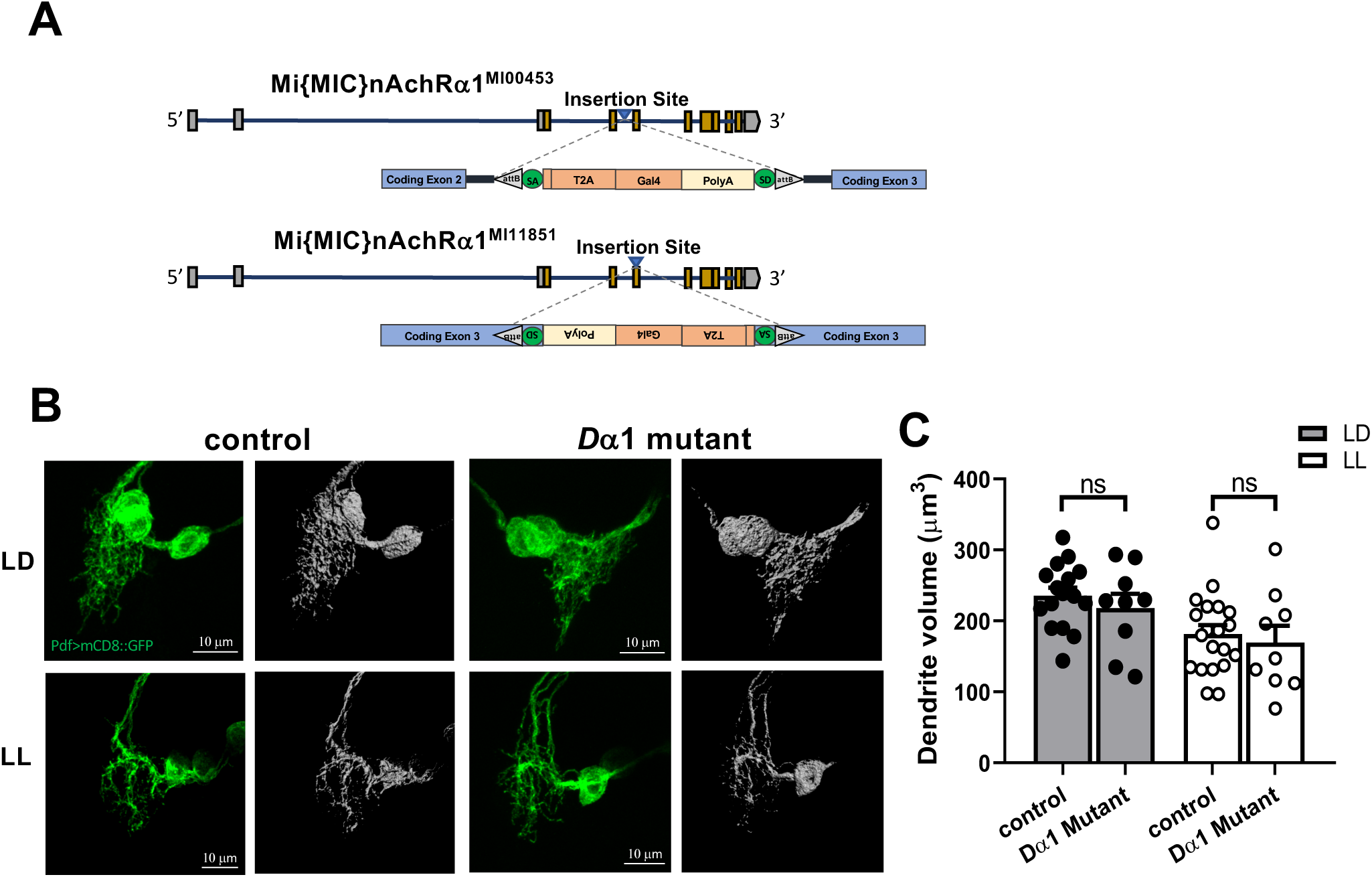
The *Dα1* subunit is not required for the development of LNv dendrites. (**A**) Schematic diagrams illustrating the two loss-of-function alleles of *Dα1* used in the phenotypic analysis. The intronic MiMIC line *Dα1*^*MI00453*^ is crossed with the exonic MiMIC insertion *Dα1*^*MI11851*^ and generates trans-heterozygous mutant larvae. (**B**) There are no observable dendrite development defects in the *Dα1* mutant. Representative projected confocal images of LNv dendrites labeled by mCD8::GFP (left, green) and their 3D reconstructions (right, grey) are shown. The light conditions and genotypes are as indicated. (**C**) Quantification of the LNv dendrite volume indicating no differences between the control group and the *Dα1* mutants. Sample size n represents the number of larvae tested. n=9-20. Statistical significance is assessed by one-way ANOVA with Tukey’s post hoc test. ns: not significant. Error bars represent mean ± SEM.

**Supplementary Figure 3.**
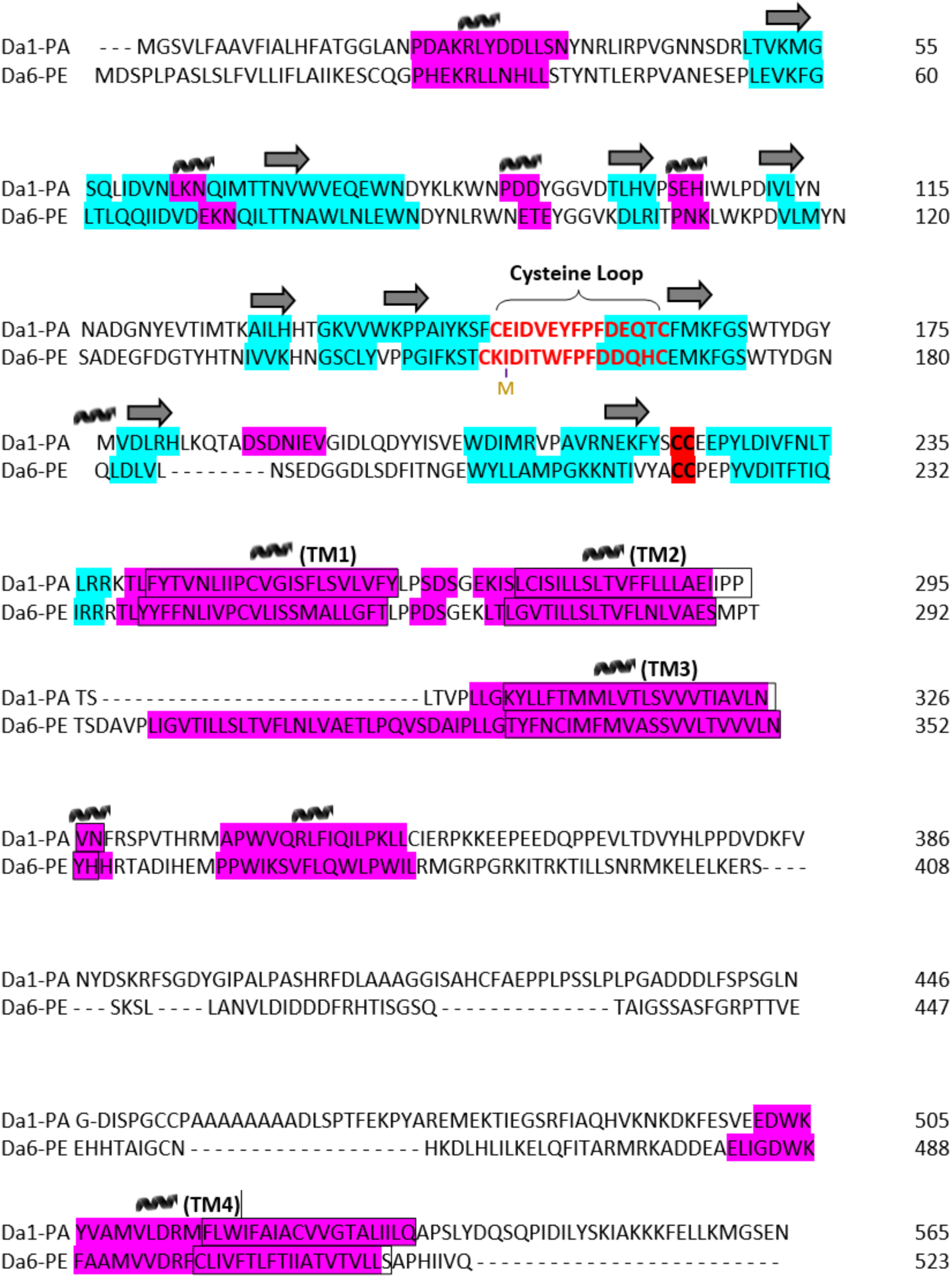
Sequence alignment of the *Dα1* and *Dα6* coding regions identifies similarities and differences at key structural motifs. Alignment of the *Dα1* and *Dα6* amino acid sequences by Clustal Omega is shown. Both subunits share the characteristics of Cys-loop ligand-gated ion channels, including the 15 residue Cysteine loop and the four transmembrane domains. The majority of differences between the subunits are found in the the secondary structures of the extracellular domain as well as the length of the TM3-TM4 intracellular loop. Additionally, variation at six residues within the Cysteine loop, plus an RNA A→I editing event occurring at I156, could lead to functional consequences. Purple: α-helix; Blue: β-Sheet; Boxed motifs: Transmembrane domains (TM1-4); Cysteine loop and C loop vicinal cysteines are colored and highlighted in red, respectively.

